# Structure-function characterization of the conserved regulatory mechanism of the *Escherichia coli* M48-metalloprotease BepA

**DOI:** 10.1101/2020.07.30.230011

**Authors:** Jack A. Bryant, Ian T. Cadby, Zhi-Soon Chong, Yanina R. Sevastsyanovich, Faye C. Morris, Adam F. Cunningham, George Kritikos, Richard W. Meek, Manuel Banzhaf, Shu-Sin Chng, Andrew L. Lovering, Ian R. Henderson

## Abstract

The asymmetric Gram-negative outer membrane (OM) is the first line of defence for bacteria against environmental insults and attack by antimicrobials. The key component of the OM is lipopolysaccharide, which is transported to the surface by the essential lipopolysaccharide transport (Lpt) system. Correct folding of the Lpt system component LptD is regulated by a periplasmic metalloprotease, BepA. Here we present the crystal structure of BepA from *Escherichia coli,* solved to a resolution of 2.18 Å, in which the M48 protease active site is occluded by an active site plug. Informed by our structure, we demonstrate that free movement of the active site plug is essential for BepA function, suggesting that the protein is auto-regulated by the active site plug, which is conserved throughout the M48 metalloprotease family. Targeted mutagenesis of conserved residues reveals that the negative pocket and the TPR cavity are required for function and degradation of the BAM complex component BamA under conditions of stress. Lastly, we show that loss of BepA causes disruption of OM lipid asymmetry, leading to surface exposed phospholipid.

**Importance:** M48 metalloproteases are widely distributed in all domains of life. *E. coli* possesses four members of this family located in multiple cellular compartments. The functions of these proteases are not well understood. Recent investigations revealed that one family member, BepA, has an important role in the maturation of a central component of the LPS biogenesis machinery. Here we present the structure of BepA and the results of a structure guided mutagenesis strategy, which reveal the key residues required for activity.

## Introduction

The outer membrane (OM) of Gram-negative bacteria is the first line of defence against environmental insults, such as antimicrobial compounds (1, 2). As such, the integrity of the OM must be maintained lest the bacteria become susceptible to stresses to which they would otherwise be resistant. The OM consists of an asymmetric bilayer of phospholipids and lipopolysaccharide (LPS) decorated with integral outer membrane proteins (OMPs) and peripheral lipoproteins. The impermeable nature of the OM can be attributed to several characteristics of the LPS leaflet, such as dense acyl chain packing, intermolecular bridging interactions and the presence of O-antigen carbohydrate chains (1, 3–7).

All the components required to construct the OM are synthesized in the cytoplasm. Specialized systems transport these molecules across the cell envelope and assemble the molecules into the OM in a coordinated fashion. Central to this is the β-barrel assembly machinery (BAM) complex. In *Escherichia coli,* the BAM complex is composed of two essential subunits, the OM β-barrel BamA and the lipoprotein BamD, and three non-essential accessory lipoproteins, BamB, BamC and BamE (8–11). The BAM complex is responsible for assembly of the Lpt system, which traffics LPS from the cytoplasm to the outer leaflet of the OM in order to maintain OM permeability barrier function (12–14). The Lpt machinery is comprised of three modules: the IM localized LptBFGC complex, which flips the LPS molecule across the IM and energizes the system; LptA, which forms a bridge between the IM and OM along which the LPS travels; and the OM complex LptD/E (12, 15, 16). The C-terminus of LptD forms an OM β-barrel which facilitates insertion of LPS directly into the outer leaflet of the OM (17, 18). The N-terminus encodes a periplasmic domain that interacts with the LptA bridging molecule (19). The two LptD domains are connected via two disulphide bonds, at least one of which is required for efficient function of the LptD/E complex (20, 21). Formation of the correct LptD disulphide bonds is reliant upon the periplasmic thiol-disulphide oxidoreductase, DsbA, as well as proper folding and insertion of the LptD β-barrel into the OM. The latter step is dependent on the BAM complex, and the interaction of LptD with its cognate OM lipoprotein partner LptE (20, 21). To be effective at LPS delivery and to maintain the integrity of the OM, maturation of the LptD/E complex is tightly regulated. The proteases DegP, BepA and YcaL each have specific roles in LptD maturation. DegP is responsible for the degradation of misfolded LptD in the periplasm, whereas YcaL targets LptD which has docked with the Bam machinery, but stalled at an early step in folding. Lastly, BepA degrades LptD which has engaged with the Bam machinery but stalled during insertion of a nearly complete barrel (22).

Amongst the LptD quality control proteases, BepA is different in that it also has chaperone activities, influences the insertion of other OMPs into the outer membrane and its deletion renders cells sensitive to multiple antibiotics (23, 24). The primary sequence of BepA indicates that this protein is a member of the M48 family of zinc metalloproteases, of which there are four in *E. coli.* The M48 proteases are characterized by an HExxH motif on the active site helix (25). The histidine residues within this motif act, usually with a third amino acid and a water molecule, to coordinate the metal ion, typically zinc, at the active site (26–28). In addition to the N-terminal M48 protease domain, BepA has a C-terminal tetratricopeptide repeat (TPR) domain. TPR domains consist of a number of stacked repeats of α-helix pairs, together forming a solenoid-like structure that is known to facilitate protein-protein interactions and multiprotein complex formation (29). Narita *et al.* reported that BepA has a dual role, degrading misfolded LptD, but also promoting correct folding and accumulation of the mature disulphide isomer of LptD (23). Further to this, the BepA protease has been shown to interact with the main BAM complex component, BamA, and to degrade BamA under conditions of stress created by the absence of the periplasmic OMP chaperone SurA (23).

Following the work of Narita *et al.*(23) we sought to determine the structure of BepA to understand the roles of the TPR and M48 peptidase domains in substrate recognition and processing. During this study two papers from other groups were published using similar structural approaches. First, Daimon *et al*.(30) determined the crystal structure of the TPR domain of BepA in isolation and observed that this domain presents a negatively charged face which was postulated to recognize components of the Bam complex and LptD. Using protein cross-linking analysis, residues of the TPR domain were demonstrated to interact with BamA, BamC, BamD and LptD. Mutation of the TPR residue F404 resulted in decreased proteolysis of BamA indicating that this residue is involved in targeting of the M48 protease domain of BepA to this substrate. More recently, Sharizal *et al.*(31) presented a full-length structure of the BepA TPR and M48 protease domains solved to 2.6 Å. In this structure the negatively charged TPR face noted by Daimon *et al*.(30) is largely buried, forming a peripheral association with the M48 protease domain. Using SAXS and engineered disulphide bonds, the potential for the TPR domain and M48 domain to move relative to one another was explored but the cross-linking experiments demonstrate that the TPR and M48 domains likely remain in tight association. Whilst multiple mutations were made, designed from the full-length structure of BepA, none of these mutations lead to any significant observable phenotype when expressed in *E. coli.*

Here we present our independently solved 2.19 Å structure of near full-length BepA, encompassing the TPR and M48 domains. Our structure largely agrees with that of Sharizal *et al.*(31), providing further evidence that TPR movement relative to the M48 domain is unlikely to be a mechanism of BepA function. Additionally, we noted the presence of an active site plug, the TPR cavity and the negatively charged pocket formed by the association of the BepA TPR and M48 domains, which we targeted for further study. Using structure-led mutagenesis studies we probed the roles of these three BepA structural elements and identify key residues in each that are required for BepA function. Furthermore, the active site plug of BepA is a structural element conserved in the M48 protease family and so our findings have broad ramifications for proteases involved in processing varied substrates across all domains of life.

## Results

### The BepA structure reveals a nautilus-like structure with TPR:protease contacts

The crystal structure of BepA_L44-Y484_ was solved to a resolution of 2.18 Å by experimental phasing using the endogenous zinc co-purified with recombinant BepA protein, present in our structure at a 1:1 stoichiometry with BepA (data collection and refinement statistics reported in Table 1); we observe a single copy of BepA in the asymmetric unit. The structure revealed the TPR domain, consisting of 12 α-helices forming 4 TPR motifs and four non-TPR helices, in tight association with the M48 zinc-metallopeptidase domain. This forms a nautilus-like fold with the TPR subdomain cupping the metallopeptidase domain **(Fig 1)**. The high-resolution data presented here are in broad agreement with that presented previously (30, 31), however there are some differences of note. The BepA TPR sub-domain was previously annotated as being composed of four TPR motif helix pairs and two non-TPR helices (nTH1 and nTH2), we have thus adopted this nomenclature.

**Figure 1.**
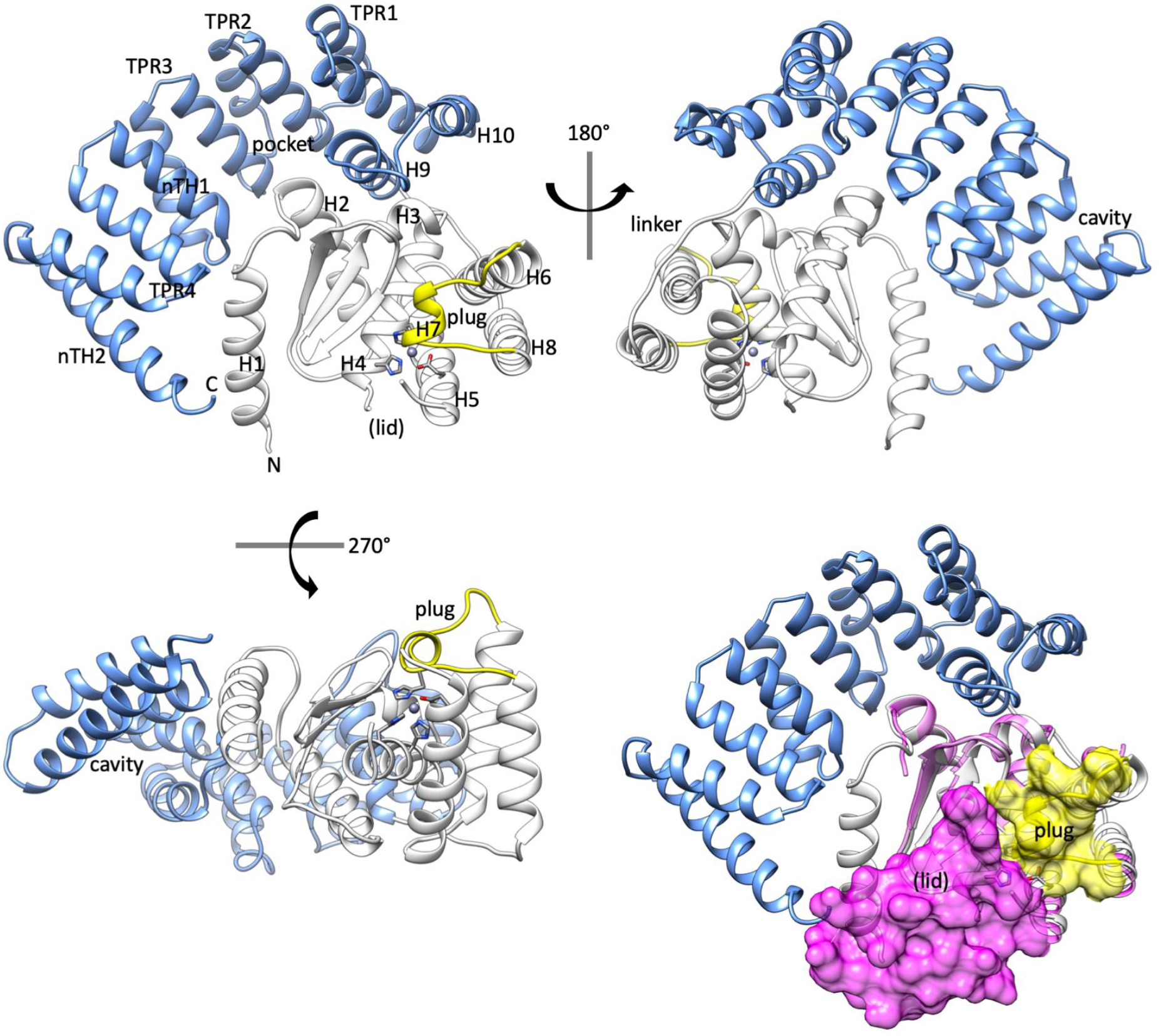
The structure of BepA reveals an occluded active site. Cartoon schematic of the X-ray crystallography structure of BepA, solved to a resolution of 2.18 Å. The TPR domain is represented in blue and the protease domain in white with the active site plug in yellow. Also labelled are the N- and C-termini, TPR pocket, TPR cavity, the linker and the site at which we expect the active site lid (lid). Important active site residues H136, H140, H246 and E201 are shown by stick diagram. The TPR motifs 1-4, non-TPR helices 1 and 1 (nTH1 and nTH2), helices, sheets and the plug labelled. Alignment of the structure presented here with that of the *G. sulfurreducens* M48 metalloprotease (PDB: 3C37 – Magenta ribbon) reveals occlusion of the active site by the potential active site lid (lid) represented as magenta surface density from the 3C37 structure and the active site plug (plug) represented as yellow surface density.

**Table 1.** Data collection and refinement statistics

|  |  |
| --- | --- |
| **Data collection** |  |
| Space group | P 21 21 21 |
| Cell dimensions |  |
| *a*, *b*, *c*, Å | 53.12, 77.02, 124.60 |
| $\alpha$, $\beta$, $\gamma$, ° | 90.00, 90.00, 90.00 |
| Resolution, Å | 77.02 - 2.18 |
| *I* / $\sigma I$ | 2.65 (at 2.18 Å) |
| Completeness, % | 98.9 (91.5) |
| Redundancy | 16.5 (6.0) |
| **Refinement** |  |
| Resolution, Å | 2.18 |
| No. of reflections |  |
| Rwork/Rfree | 0.176 / 0.201 |
| No. of atoms | 3269 |
| Protein | 3143 |
| Ligand/ion | 6 |
| Water | 120 |
| *B*-factors | 47.0 |
| Protein | 43 |
| Ligand/ion | 71 (SO<sub>4</sub>), 37 (Zn) |
| Water |  |
| rmsd |  |
| Bond lengths, Å | 0.006 |
| Bond angles, ° | 1.00 |

\*Values in parentheses are for highest-resolution shell.

Our structure demonstrates that the TPR domain consists of 12 α-helices, whereas the structure was previously annotated with 10 α-helices in order to maintain nomenclature with the previously solved TPR-domain structure of residues 310-482 (30, 31). Despite the previous annotation as non-TPR helices, we observe that helices 8 and 9 form part of the TPR domain and are preceded by an extended linker region, residues M263-S271, which connects the TPR domain to helix 7 of the protease domain **(Fig 1).** Helices 8 and 9 contribute a tight turn at the end of the TPR domain, allowing the M48 metallopeptidase domain to be cupped by the pocket formed from TPR motifs 2 and 3 **(Fig 1).** Interaction of the protease domain with the TPR pocket creates a larger negatively charged pocket, which is also noted in the structure presented by Shahrizal *et al*. (31). The context provided by the full-length protein structure shows that while the TPR pocket interacts with the protease domain, the TPR cavity is positioned away from the protease active site on the opposite side **(Fig 1).** The cavity also comes into close proximity with the N-terminal helix, which is contained within the protease domain, therefore potentially facilitating TPR:protease domain communication.

The protease domain of BepA consists of the active site α-helix H4 containing the HExxH motif, and an active site plug formed by a loop between helices H6 and H7, residues S246-P249. We did not observe any density corresponding to positions L146-I194 and considering that this section is in close proximity to the active site, we expect that it may form a dynamic regulatory region **(Fig 1).** We sought to find evidence that the unresolved area of the protein may correspond to a dynamic lid. Therefore, we scrutinized the Protein Data Bank (PDB) for similar structures. Information on the missing region of our structure can be inferred from an unpublished structure in the PDB of an M48 zinc-metallopeptidase from *Geobacter sulfurreducens,* which consists of only the protease domain, with no associated TPR (PDB: 3C37). The structure of the *G. sulfurreducens* protease structure provides some information on the missing section and demonstrates a short three-turn extension to the C-terminus of active site helix H4, beyond that seen in the BepA structure. This is followed by a glycine facilitated kink and another three helical turns terminating at residue D136 of the 3C37 structure (**Supplementary Fig S1**). The 3C37 structure is also missing a section, D136-N139, however residues M140-F149 form another short α-helical region, which is connected to the N-terminus of helix H5, by an extended region formed by residues G150-S158 of the 3C37 structure **(Supplementary Fig S1).** While also incomplete, the recently published BepA structure also provides some information on this section, which is also largely in agreement with that of the 3C37 structure **(Supplementary Fig S1)** (31). Overall, comparison of the structure presented here, that of Shahrizal *et al*. (31) (PDB: 6AIT), and the *G. sulfurreducens* structure (PDB: 3C37), suggests that the missing section from the structure presented here may form a putative active site lid. The putative lid, along with the plug, likely regulates access to the active site as alignment of the three structures shows that the lid and plug occlude the active site **(Fig 1)**. The fact that no density for the putative lid is observed in our structure, and that partial sections are missing in those presented previously, suggests that the active site lid is dynamic and may adopt multiple conformations.

### Mobility of the conserved active site plug is required for BepA function

The structure shows the HExxH motif, which is characteristic of zinc-dependent metallopeptidases (25, 27) and is found within helix H4 **(Fig 1)**. The active site zinc ion is coordinated by H136 and H140 within the HExxH motif, E201 contributed by helix H5, and H246 on a loop that forms the small α-helical active site plug **(Fig 1)**. Multiple alignment of the four *E. coli* M48 metallopeptidases, HtpX, YcaL, LoiP and BepA demonstrates that zinc coordinating residues are all conserved along with the proline following H246, P247, and an arginine further towards the C-terminus, R252, which resides within the active site **(Fig 2A and Fig 3A).** Analysis of the HMM logo generated for the M48 metallopeptidase family demonstrated that not only is the HExxH motif and the zinc-coordinating glutamic acid conserved, but the H-P-x(4)-R motif within the active site plug is also conserved throughout the whole pfam family (PFO1435), which includes proteins from all domains of life (**Fig 2B)**. The active site zinc ion is usually chelated by three amino acid residues and one water molecule, which is utilized to catalyze proteolysis of the substrate (26, 28). Co-ordination of the zinc ion by H246 fulfils the fourth ligand, therefore suggesting that a rearrangement of the active site plug should be required for proteolytic activity. Alignment with the structure of human nuclear membrane zinc metalloprotease, ZMPSTE24, with a bound substrate peptide (PDB: 2YPT) reveals that the BepA active site plug occupies the same physical space as the substrate for ZMPSTE24 would occupy **(Fig 3B).** Residue H246 on the BepA active site plug directly clashes with positioning of substrate in the 2YPT structure and the hydrophobic residues I242 and L243 occupy a similar space to the 2YPT substrate hydrophobic residues I3’ and M4’ **(Fig 3B)**. Based on these observations, we hypothesized that the active site plug occludes the active site and is likely to relocate in order to facilitate substrate access to the active site **(Fig 1 and Fig 3B)**.

**Figure 2.**
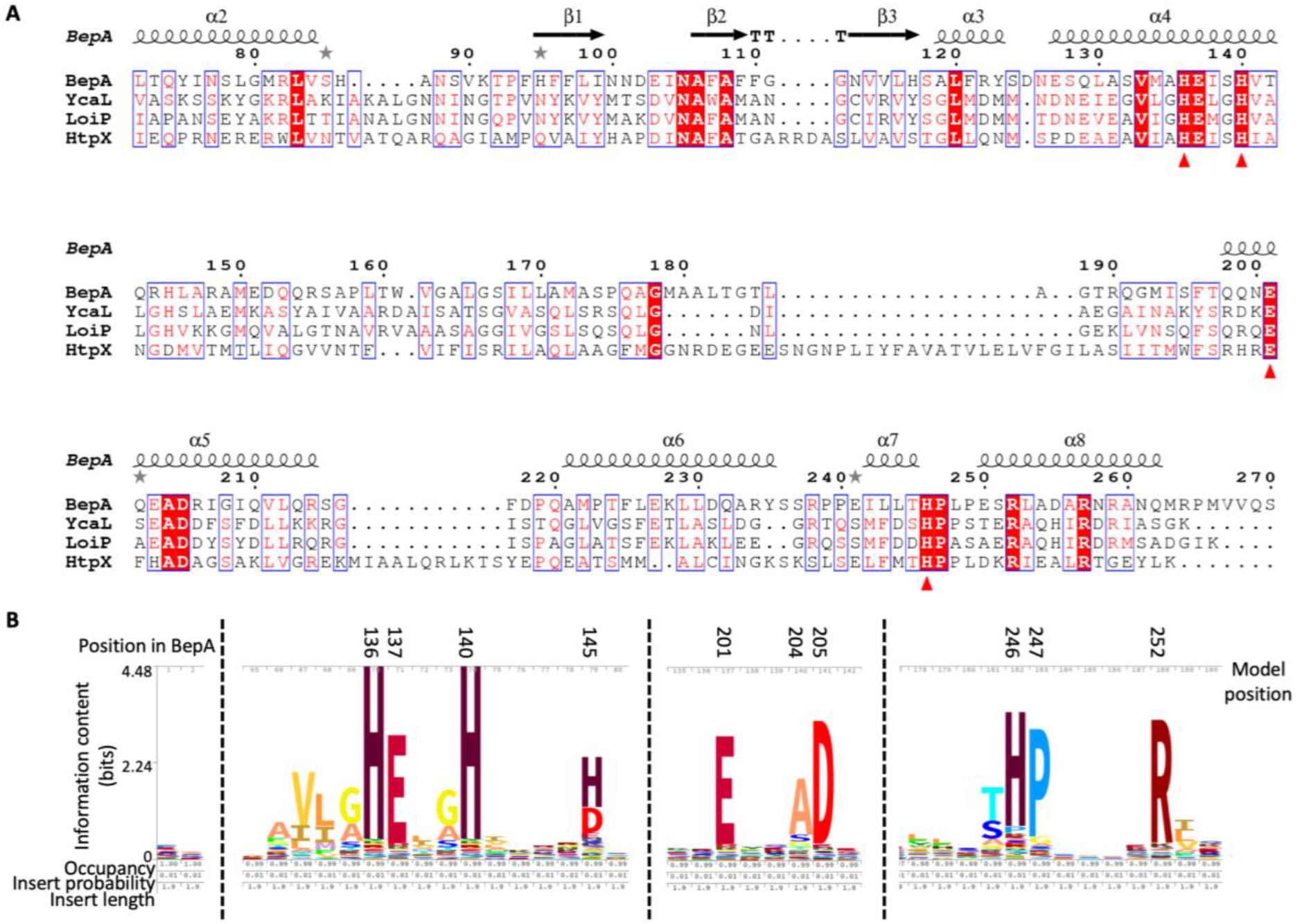
Conservation of the M48 metalloprotease HExxH motif and active site plug residues. **A.** Amino acid sequences for *E. coli* BepA, YcaL, LoiP and HtpX were submitted to Clustal Omega (https://www.ebi.ac.uk/Tools/msa/clustalo/) in order to generate a multiple alignment to allow analysis of amino acid conservation and subsequently processed using ESPript 3.0 (http://espript.ibcp.fr) (49, 50). Sequences are named on the left and numbered above the line based on the BepA sequence. Gaps in the alignment are represented by dots. A single fully conserved residue is highlighted red and the zinc co-ordinating residues are labelled with a red triangle under the residue. BepA secondary structure is represented on the top line with α-helices labelled with a spiral and β-sheets by arrows. The protease domain sub-section of the alignment is shown, for the full alignment see Figure S3. **B.** HMM logo generated for the pfam M48 family of metalloproteases (PF01435) from the pfam website (https://pfam.xfam.org) with the HMM profile constructed on the pfam seed alignment. Three sections of the alignment are shown, which demonstrate conservation of the active site zinc co-ordinating residues H136, H140, E201 and H246. The active site plug clearly contains a conserved motif, H-P-x(4)-R. Amino acid position in BepA and within the model are both shown with the stack height corresponding to information content (bits), which represents the invariance of the position. Letter height divides the stack height according to letter frequency.

To test the importance of H246 in occupying the fourth coordination site on the zinc ion, we generated a mutation of the H246 position to asparagine (H246N). For comparison, we also constructed the E137Q mutation in the active helix HExxH motif, which has previously been shown to prevent protease activity of BepA (23). To test whether the H246N BepA mutant is functional, we assayed the ability of this mutant to complement the Δ*bepA* strain, which is known to exhibit increased sensitivity to large antibiotics such as vancomycin, presumably due to impaired barrier function of the OM. The E137Q active site mutant was incapable of restoring vancomycin resistance to Δ*bepA* cells and had a severe negative effect on the growth of the Δ*bepA* mutant **(Fig 3C)**. The H246N mutant BepA was also incapable of complementing vancomycin sensitivity of the Δ*bepA* cells; however, while the H246N protein also severely increased the vancomycin sensitivity of the mutant beyond that of the empty vector control, the negative effect was less extreme than with the E137Q version of the protein **(Fig 3C)**. Considering this phenotype, we decided to investigate if the mutated proteins had a dominant-negative effect in the parent background expressing wildtype *bepA*. We found that the empty vector and wild-type BepA had no detrimental effect on BW25113 parent cells grown in the presence of vancomycin. Our analysis of the E137Q mutant was in agreement with previous studies when analyzed in the parent background and demonstrated a severe dominant-negative phenotype (23). We also observed that the presence of H246N BepA had a dominant-negative effect on the capacity of the cells to grow in the presence of vancomycin, despite the presence of wild-type BepA expressed from the chromosomal locus. Similar to the effect in the mutant background, the dominant-negative effect of the H246N protein was less severe than that of the E137Q derivative **(Fig 3C)**. We speculate that this is may be because the active site plug is less able to interact with the active site zinc ion and that the protein may be in a constitutively activated or “de-regulated” conformation (**Fig 1 and Fig 3**). Western blotting to detect the expression of BepA proteins in whole cell lysates using anti-6xHis antibodies showed an elevated level of the E137Q protein compared to wild-type and an absence of observable tagged protein in the H246N sample. These observations were consistent between the *ΔbepA* and parent backgrounds (**Supplementary Fig S2**). These results support the hypothesis that the E137Q mutation renders BepA protease inactive, therefore stabilizing the protein due to a lack of auto-proteolytic activity, which has been observed previously (23). Considering that the H246N BepA has a dominant-negative effect, the absence of a detectable tagged protein by western blot suggests the C-terminal His-tag may be auto-proteolytically degraded, an observation that has previously been made for the wild type BepA protein (23). This data supports the hypothesis that the H246N mutation gives rise to a protein with de-regulated proteolytic activity.

**Figure 3.**
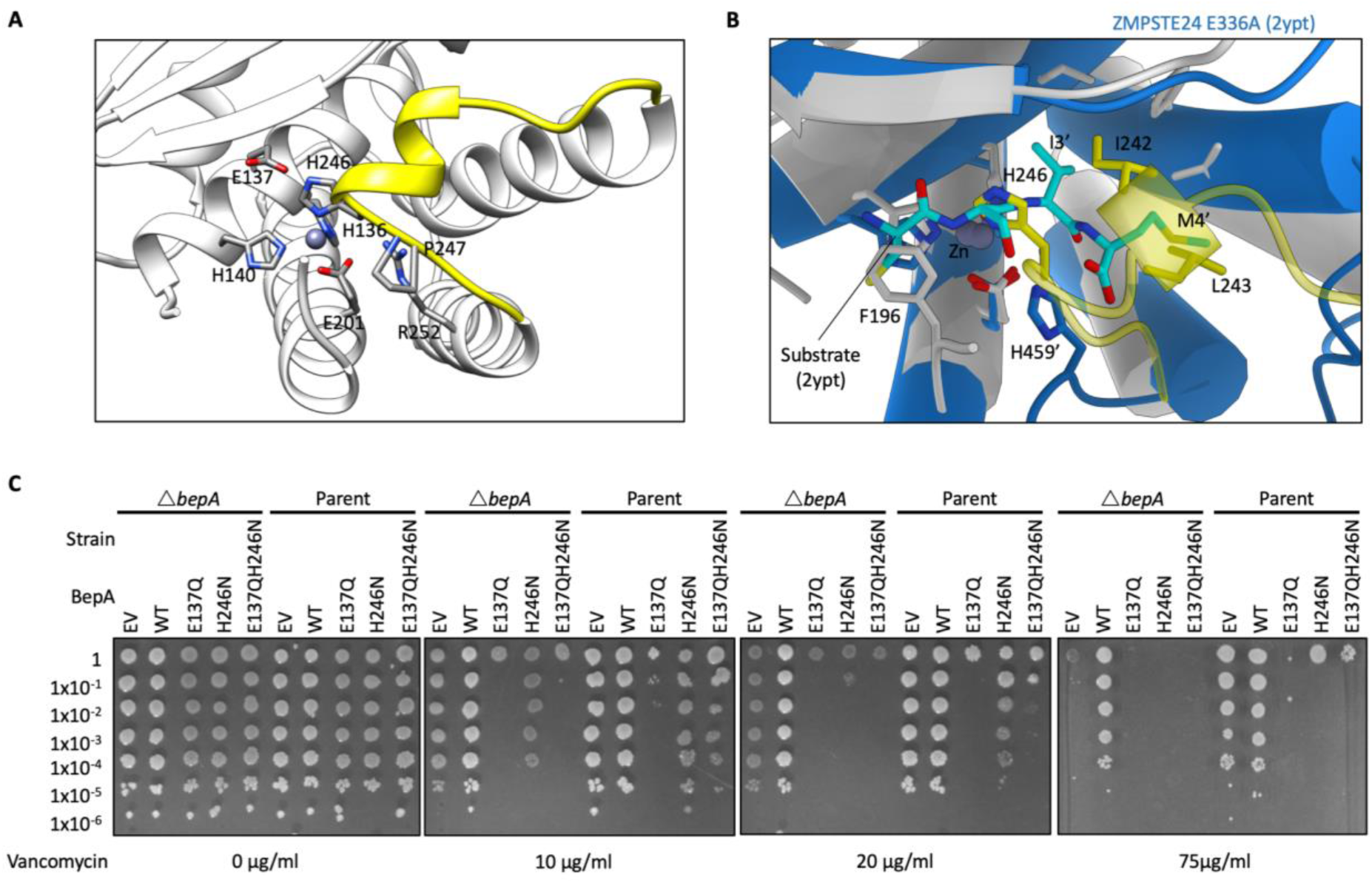
The BepA active site plug acts to regulate BepA proteolytic activity. Analysis of the BepA structure suggested a regulatory role for the active site plug, therefore plasmids encoding mutated *bepA* were screened for their capacity to complement the vancomycin sensitivity of Δ*bepA E. coli.* **A.** Structural diagram of the BepA active site with key conserved residues represented by stick diagram and labelling. **B.** Alignment of the BepA active site (transparent white and yellow ribbon) with that of the human nuclear membrane zinc metalloprotease ZMPSTE24 mutant E336A with a synthetic substrate peptide (PDB: 2YPT) (51) (Opaque blue ribbon) 2YPT residues are labelled with the addition of a ‘ symbol. **C.** Screen for vancomycin sensitivity of cells carrying pET20b encoding WT or mutated copies of BepA in the parent or Δ*bepA* strain background. The empty vector control is labelled EV. Cells are normalised to OD_600_ = 1 and ten-fold serially diluted before being spotted on the LB agar containing the indicated antibiotics (all plates contain 100 μg/ml carbenicillin additionally).

In order to test if the auto-proteolysis of the H246N protein was due to increased protease activity, we introduced the established protease dead mutation E137Q. The BepA E137Q H246N substitution was not capable of complementing the vancomycin sensitivity and had a severe dominant-negative effect similar to that of E137Q alone **(Fig 3C)**. Analysis of the E137Q H246N BepA protein by western blot showed a similar level of tagged protein to the E137Q protein **(Supplementary Fig S2)**. These data suggest that introduction of the E137Q mutation either prevents auto-proteolysis of the C-terminal 6xHIS tag in the H246N mutant or alternatively stabilizes the protein, preventing it from being targeted by other periplasmic proteases.

The importance of residue H246 for BepA function, and the conformation of the active site plug observed in our crystal structure, suggests this may be an inactive form of the protein. Therefore, we hypothesized that movement of the active site plug must be required to facilitate substrate access to the active site. We aimed to tether the active site in the conformation observed in our crystal structure by engineering a disulphide bond. Cysteine substitutions were introduced into proximal sites in BepA, specifically at positions E103, in the loop between S1 and S2, and E241 in the active site plug, either individually or in concert **(Fig 4A)**. The single cysteine substitutions complemented the sensitivity phenotype, indicating that the single substitutions had no impact on BepA function. However, the double cysteine mutant was incapable of restoring vancomycin resistance to the *bepA* mutant under normal growth conditions. In contrast, in the presence of the reducing agent TCEP (tris(2-carboxyethyl)phosphine), the double cysteine mutant was able to complement vancomycin sensitivity **(Fig 4B)**. The double cysteine mutant also caused a severe dominant negative effect in the parent background, which was alleviated by the presence of the reducing agent TCEP, therefore allowing free movement of the regulatory active site plug and normal functioning of BepA **(Fig 4B).** These observations suggest that in the E103C E241C BepA a disulphide bond was formed that tethered the active site plug in the inactive conformation, causing similar effects to the E137Q protease-dead mutation and that free movement of the plug is essential to function **(Fig 4B)**.

**Figure 4.**
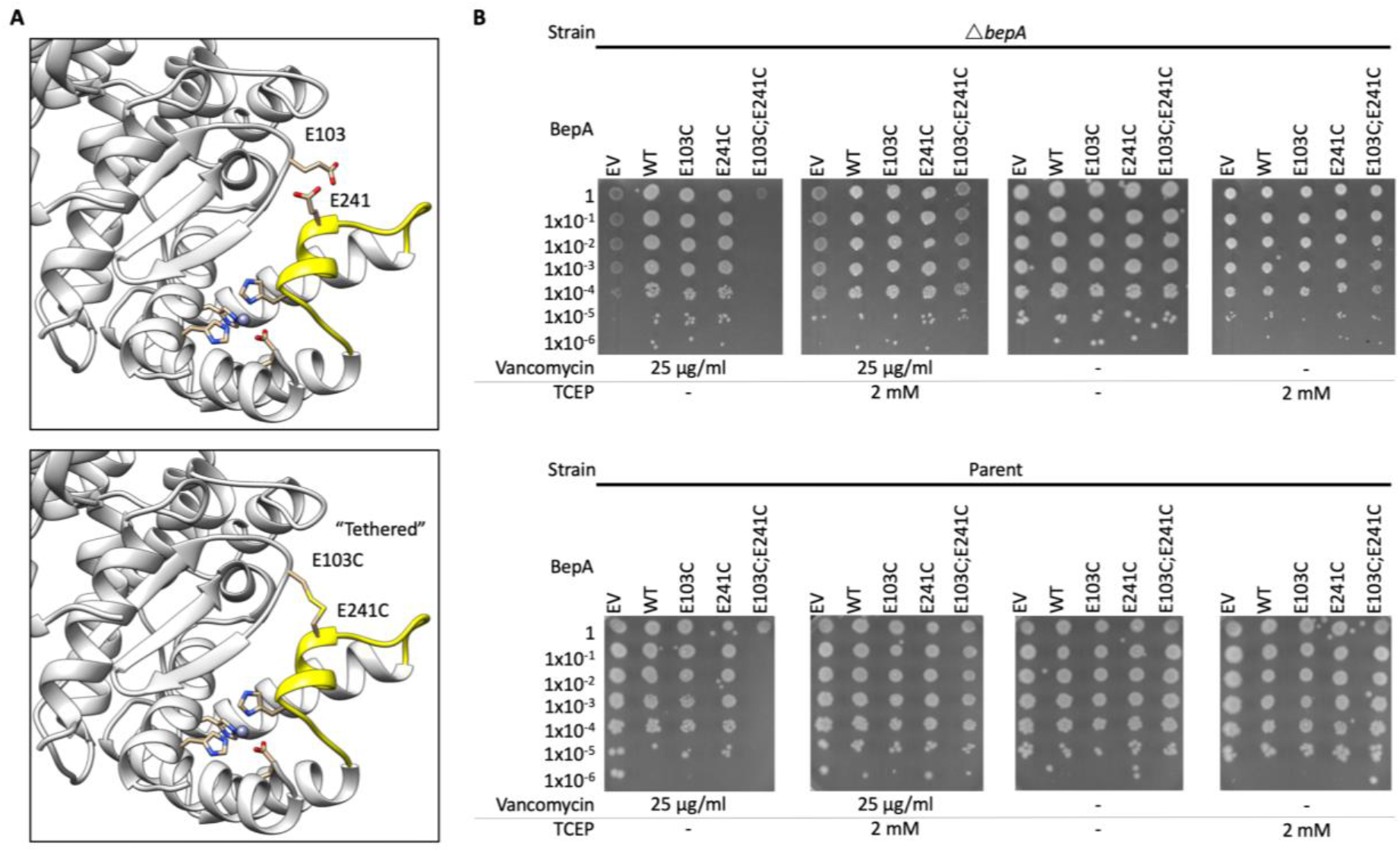
Flexibility of the active site plug is required for BepA function. The requirement for flexibility of the BepA active site plug for full BepA function was assayed by disulphide bond tethering of the active site plug and functional screening. **A.** Structural representation of the BepA active site with residues targeted for mutation to cysteine, E103 and E241, labelled and coloured yellow. **C.** Screen for vancomycin sensitivity of cells carrying pET20b encoding WT or mutated copies of BepA in the parent or Δ*bepA* strain background. The empty vector control is labelled EV. Cells are normalised to OD_600_ = 1 and ten-fold serially diluted before being spotted on the LB agar containing the indicated antibiotics or reducing agent TCEP (all plates contain 100 μg/ml carbenicillin additionally).

### The BepA negative pocket and TPR cavity are required for function and BepA-mediated degradation of BamA

The TPR domain contains two potential substrate binding sites, termed the “pocket” on the protease proximal face and the “cavity” on the protease distal face **(Fig 1 and Fig 5)**. We identified two conserved charged residues, R280 and D347, in the BepA TPR pocket, which forms a larger negatively charged cleft through interaction with the protease domain **(Fig 5A)**. The negatively charged cleft is connected to the active site via a negatively charged ditch and has previously been hypothesized to facilitate substrate interactions (31). However, no evidence for the importance of this site for BepA function has yet been provided. We targeted these two conserved charged residues within the pocket, and vancomycin sensitivity assays revealed that the R280 mutations had no significant effect on complementation of the *bepA* mutant. However, the D347R mutation had a mild negative effect on the capacity of the BepA protein to complement the vancomycin sensitivity of the *bepA* mutant and a dominant-negative effect in the parent background, despite the protein being expressed to a lower level than the WT protein **(Supplementary Fig S2 and S3)**.

**Figure 5.**
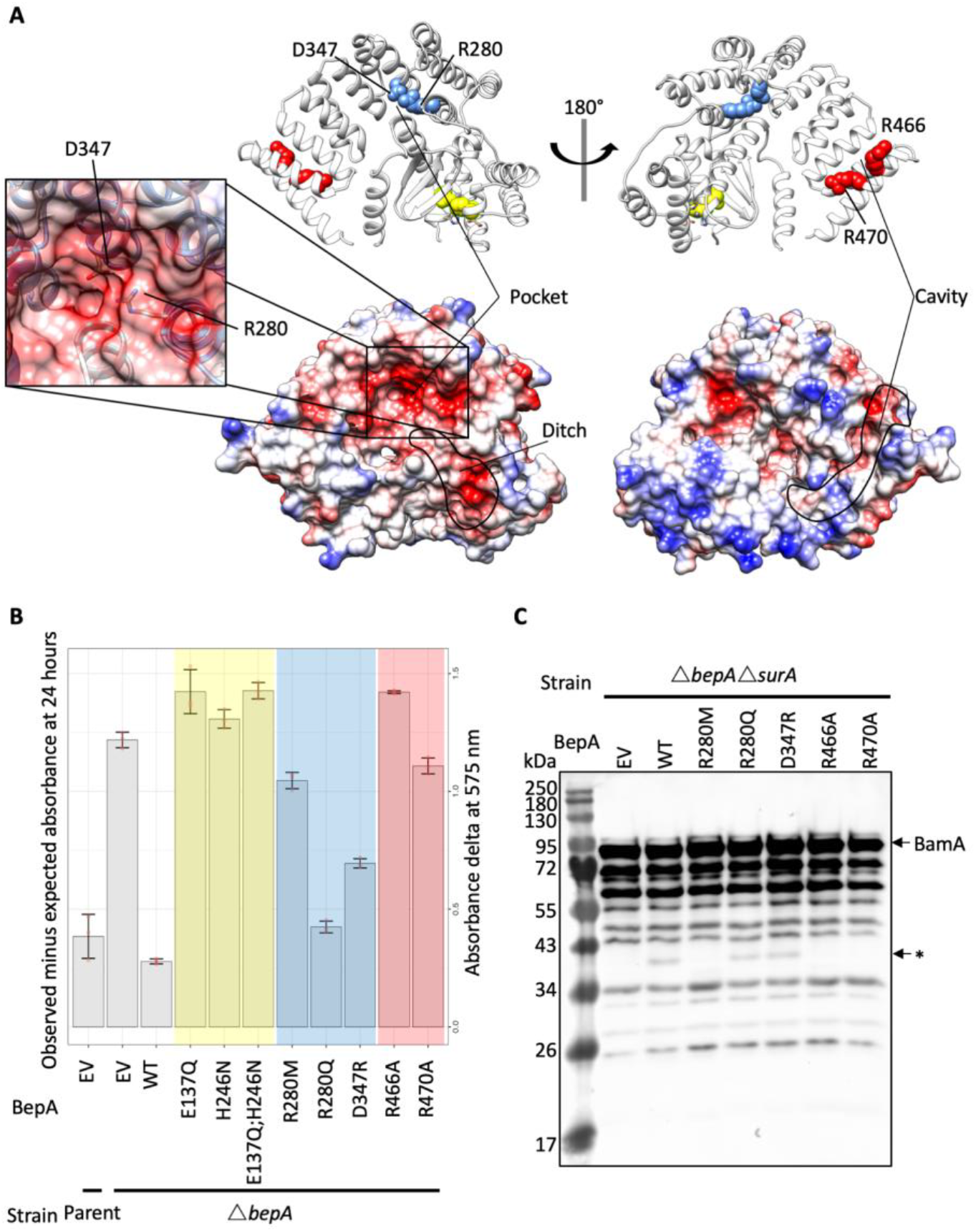
Conserved residues in the pocket and TPR cavity are required for function. The BepA pocket and TPR cavity contain conserved residues that were mutated to test their importance for BepA function. **A.** Structure of BepA showing residues targeted for mutation as colour-coded spheres that match the colour-coded chart represented in panel B. Also shown are surface representations of BepA in the same orientations coloured according to surface charge from red for negatively charged, through white for near neutral to blue for positive charge. The zoom-in box depicts the position of the key D347 and R280 residues. **B.** CPRG permeability assay of Parent or Δ*bepA* cells carrying pET20b with WT or mutant copies of BepA as indicated. The empty vector control is labelled as EV. The CPRG turnover score (change in absorbance at OD575 compared to the Lac^-^ cells) is represented for two independent experiments each containing three replicates **C.** The R280M, R466A and R470A substitutions prevent BepA-dependent generation of a putative BamA degradation product in the Δ*surA* background. Total cellular protein extracts were prepared from Δ*bepAΔsurA* cells carrying pET20b encoding WT or mutated copies of BepA. The empty vector control is labelled EV. Samples were separated by SDS-PAGE and transferred to nitrocellulose membrane for western immuno-blotting using antisera raised in rabbits against the BamA POTRA domain. The putative BamA degradation product is labelled with an arrow and asterisk (*), with the full length BamA labelled.

The effect of D347R is weak by comparison with the active site mutations, therefore we utilized a more sensitive permeability assay to assess the mutation. Vancomycin is a large (1450 Da) hydrophobic antibiotic that does not normally penetrate the OM. The target for vancomycin is the abundant D-alanyl-D-alanine substrate, which is must bind in sufficient quantity to exhibit an effect on cell growth/lysis. Chlorophenyl red-β-D-galactopyranoside (CPRG) is a hydrophobic β-galactosidase substrate that is smaller (585 Da), but also fails to penetrate wild-type *E. coli.* OM permeability defects allow penetration of CPRG into the cell where it is then accessible to cytoplasmic β-galactosidase, which hydrolyses the CPRG to produce a red colour (32, 33). Production of the red colour is a sensitive indicator of cell permeability and thus can be measured using a time resolved wavescan of cells grown on LB agar supplemented with CPRG (33, 34). The BW25113 parent strain is Lac^-^, therefore strains were cotransformed with the relevant *bepA* encoding plasmid and a *lacZYA* expression vector (32, 33, 35). CPRG assays indicated that the *bepA* mutant cells are indeed more permeable to the β-galactosidase substrate CPRG and that this permeability phenotype can be complemented **(Fig 5B)**. Active site mutants E137Q or H246N that cause increased vancomycin sensitivity when compared to the *bepA* mutant empty vector control are not able to restore the OM barrier against CPRG. The conditions used for the assay here appear to be too sensitive to measure the differences between the empty vector control, E137Q and H246N mutants that are apparent from vancomycin sensitivity screening **(Fig 5B)**. However, the increased sensitivity of the assay showed that mutations altering conserved residues in the pocket (R280M, D347R) are not able to fully complement the permeability defect **(Fig 5B)**. This suggests that the phenotypes caused by these mutations are mild compared to the active site mutations. The mild permeability phenotype could explain the lack of observable vancomycin sensitivity despite increased permeability to CPRG.

We next sought to assess the cavity in the TPR domain, which has been shown to be involved in BepA binding to the Bam complex (30). Conservation analysis revealed two conserved arginine residues, R466 and R470, which have yet to be analyzed for their role in BepA function. We expected these residues might be involved in substrate recognition or interaction with protein complex partners due to the prominent position in the cavity and their high level of conservation despite any obvious structural role **(Fig 5A)**. Mutation of these residues to alanine appeared to have no impact on the capacity of the BepA protein to complement the vancomycin sensitivity phenotype **(Supplementary Fig S3)**. However, the CPRG permeability assay demonstrated that R466 and R470 are indeed required for full complementation of the OM permeability defect caused by loss of BepA **(Fig 5B)**.

BepA has been shown to degrade the BAM complex component BamA under conditions of stress induced by the absence of the chaperone SurA (23). Considering that the TPR cavity has been shown to interact with Bam complex subunits (30), and that we observed cell permeability defects on complementation with the TPR cavity mutants, we reasoned that these mutants may be defective in BepA-mediated degradation of BamA. Therefore, we analyzed whole cell lysates from Δ*bepAΔsurA* cells expressing WT BepA or the R280M, R280Q, D347R, R466A or R470A derivatives of BepA by western immuno-blotting with anti-serum raised against POTRA domain five of BamA (36). As has been demonstrated previously, we observed that introduction of WT BepA into the cells leads to generation of an anti-BamA antibody reactive BamA degradation product of approximately 40 kDa (23) **(Fig 5C)**. Production of the putative BamA degradation product was not detectable in cells expressing the R280M derivative or in cells expressing BepA with substitutions in the TPR cavity (R466A and R470A), all of which had the most severe permeability defects in this set **(Fig 5B and 5C)**. In contrast, production of the putative BamA degradation product was unaffected in cells expressing the negative pocket derivatives R280Q and D347R, which were also less permeable to CPRG than the other mutants assayed **(Fig 5B and 5C)**. These data suggest that these residues are important for BepA-mediated degradation of BamA in the absence of the chaperone SurA. This also supports the previous observation that the TPR cavity is required for interaction with the Bam complex (30).

### Loss of BepA leads to increased surface exposed phospholipid

It has been established that *bepA* mutant *E. coli* are more sensitive to hydrophobic antibiotics with a high molecular mass, such as vancomycin, erythromycin, rifampicin and novobiocin (23). This is presumed to be due to increased OM permeability. The hypothesis is that the loss of BepA results in reduced LptD assembly, therefore leading to reduced OM LPS content. This would in turn cause phospholipids to flip from the inner leaflet to the outer leaflet of the OM, creating a perturbation in OM lipid asymmetry and increased OM permeability to large antibiotics (22, 23, 37). While it has been established that the Δ*bepA* cells are more permeable, as demonstrated here by increased permeability to CPRG, this is not necessarily evidence of perturbed OM lipid asymmetry. Perturbation of OM lipid asymmetry can be detected through monitoring the activity of the enzyme PagP. On detecting surface exposed phospholipids, the OM localized Lipid A palmitoyltransferase PagP, utilizes the outer leaflet phosphoplipids as palmitate donors to convert hexa-acylated Lipid A to hepta-acylated Lipid A (38–40). The resulting lyso-phospholipid product is then degraded by the OM phospholipase PldA **(Fig 6A)**. To measure the levels of hepta-acylated Lipid A, radiolabelled Lipid A was isolated from the Δ*bepA* mutant or bacteria that had been complemented with BepA, BepA E137Q or BepA H246N. The lipids were then separated by thin layer chromatography. The parent strain BW25113, transformed with empty pET20b, were treated with EDTA prior to Lipid A isolation, a process that is known to induce high levels of hepta-acylated Lipid A production and act as a positive control (41–43). Cells lacking BepA showed a significant increase in the levels of hepta-acylated Lipid A in relation to hexa-acylated Lipid A, indicating perturbation of OM lipid asymmetry in the absence of functional BepA **(Fig 6B and 6C).** While the catalytically dead E137Q and the H246N mutants were not able to rescue this defect, they also did not appear to significantly increase the levels of hepta-acylated Lipid A compared to cells lacking BepA **(Fig 6B and 6C)**. Additionally, we did not see any effect on lipid A palmitoylation for any of the other mutations used in this study. However, we suggest that this is likely due to the sensitivity of the assay. These data demonstrate that loss of BepA leads to an increase in surface exposed phospholipid, which is likely the cause of increased permeability.

**Figure 6.**
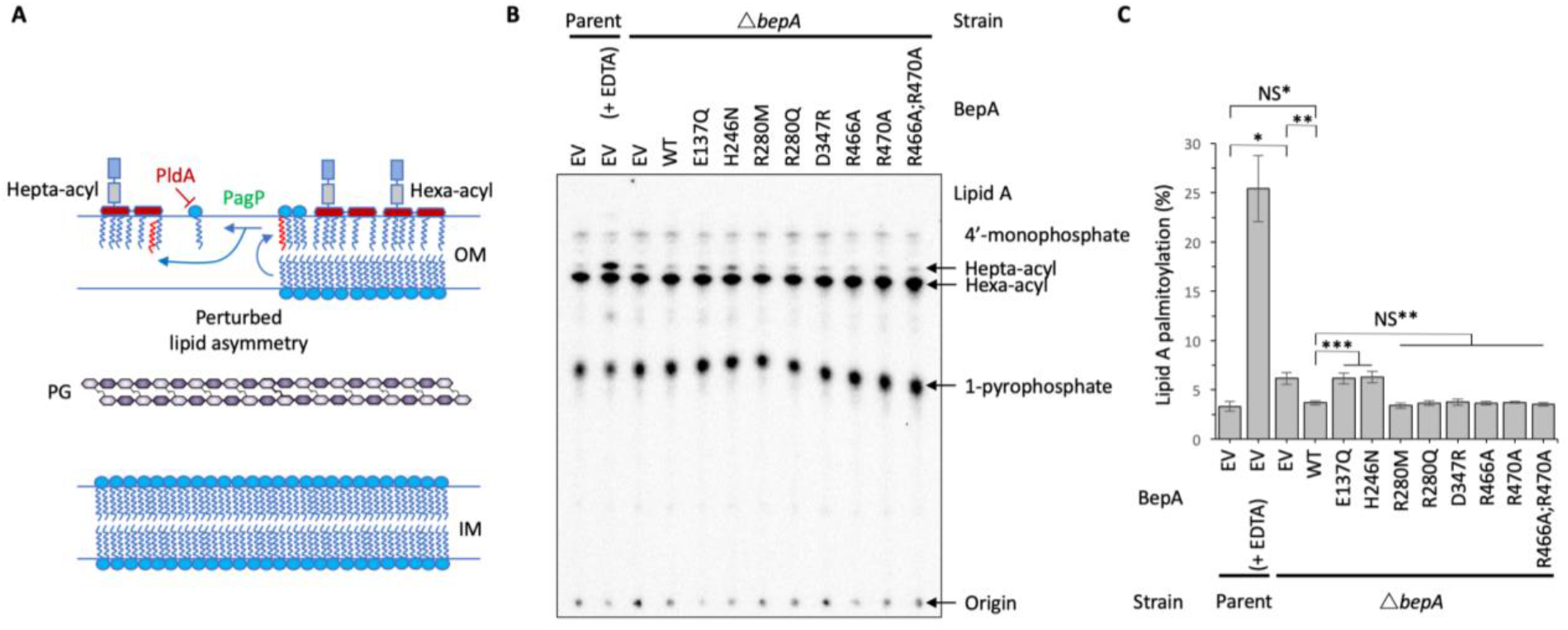
Loss of BepA leads to surface exposed phospholipid. The increased permeability of Δ*bepA* cells was hypothesised to be due to increased surface exposed phospholipid, therefore this was tested by the PagP mediated Lipid A palmitoylation assay, which detects surface exposed phospholipid. **A.** Schematic demonstrating the role of PagP in sensing and responding to surface exposed phospholipid. **B.** PagP mediated Lipid A palmitoylation assay. PagP transfers an acyl chain from surface exposed phospholipid to hexa-acylated Lipid A to form hepta-acylated Lipid A. [32-P]-labelled Lipid A was purified from cells grown to midexponential phase in LB broth with aeration. Equal amounts of radioactive material (cpm/lane) was loaded on each spot and separated by thin-layer chromatography before quantification. As a positive control, cells were exposed to 25 mM EDTA for 10 min prior to Lipid A extraction in order to chelate Mg^2+^ ions and destabilise the LPS layer, leading to high levels of Lipid A palmitoylation. **C.** Hepta-acylated and hexa-acylated lipid A was quantified and hepta-acylated Lipid A represented as a percentage of total. Triplicate experiments were utilised to calculate averages and standard deviations with students t-tests used to assess significance. Student’s *t*-tests: \**P* < 0.005 significant compared with Parent EV; \*\**P* < 0.005 significant compared with Δ*bepA* EV; *** *P* < 0.001 significant compared with Δ*bepA* WT; NS* *P* > 0.1 compared with Parent EV; NS** *P* > 0.1 compared with Δ*bepA* WT.

## Discussion

In this study, we present the structure of full-length BepA at a resolution of 2.18 Å, which is a periplasmic M48 zinc metalloprotease family protein involved in regulating the maturation of the LPS biogenesis machinery in Gram-negative bacteria. Our independently-solved structure guided our mutagenesis strategy to identify and investigate the mechanism of the BepA active site plug, which contains a conserved motif found throughout the M48-metalloprotease family. The structure presented here is missing density for a region near to the active site. Comparison to structural data available in the PDB demonstrated that this region corresponds to what appears to be an active site lid that in part occludes access to the active site residues (31). The three available structures all demonstrate some missing density within the lid, therefore this could be explained by flexibility within this region to facilitate substrate access to the active. This highlights an attractive area for future study into the regulatory mechanisms employed by BepA and the M48 metalloproteases.

In combination with the active site lid, access to the site is also blocked by the active site plug, which we focused on here. We have shown that H246N within a small helix on the active site loop coordinates the zinc in our structure and is essential for correct function of BepA. Through the use of disulphide bond tethering, we also demonstrate that the plug must be mobile for function of the protein. This suggests that the plug may act in an auto-regulatory function to either block the active site or move to facilitate access for the substrate and proteolytic activity. Three other M48 family metalloproteases are annotated in *E. coli*: the OM lipoprotein LoiP, with which BepA has been shown to interact (44); the IM heat-shock induced endopeptidase, HtpX (45); and the recently characterized OM lipoprotein YcaL, which is also involved in the regulation of LptD insertion into the OM (22). We assessed conservation within these four metalloproteases and found that the key active site plug residues are conserved amongst these proteins. Through analysis of the HMM logo for the M48 family of metallopeptidases we observed that the regulatory plug mechanism is conserved throughout the whole pfam protein family (PF01435) and is found in all domains of life (46). The regulatory plug is characterized by two conserved residues, H246 and P247, and the active site contains one further conserved arginine leading to a conserved motif, H-P-x(4)-R. Addition of the regulatory plug motif to the characteristic H-E-x-x-H-motif of zinc metallopeptidases allows the specific identification of this protein family within *E. coli* by using the online pattern search tool MOTIF2 (https://www.genome.jp/tools/motif/MOTIF2.html) using the pattern search H-E-x-x-H-x(30,140)-H-P-x(4)-R. The results of this search, in *E. coli,* identify three proteins other than the four M48 family metallopeptidases. One of these is a prophage cell-death peptidase encoded by the *lit* gene, which is classified as the single member of the U49 peptidase family. We anticipate that this peptidase and the rest of the M48 family of zinc metallopeptidases are likely to be auto-regulated by a conserved H-P-motif active site plug mechanism similar to that of BepA **(Fig 7)**. Therefore, identification of this motif will be important for identifying the active site plug mechanism in future studies involving this family of proteins, which is found in all domains of life.

**Figure 7.**
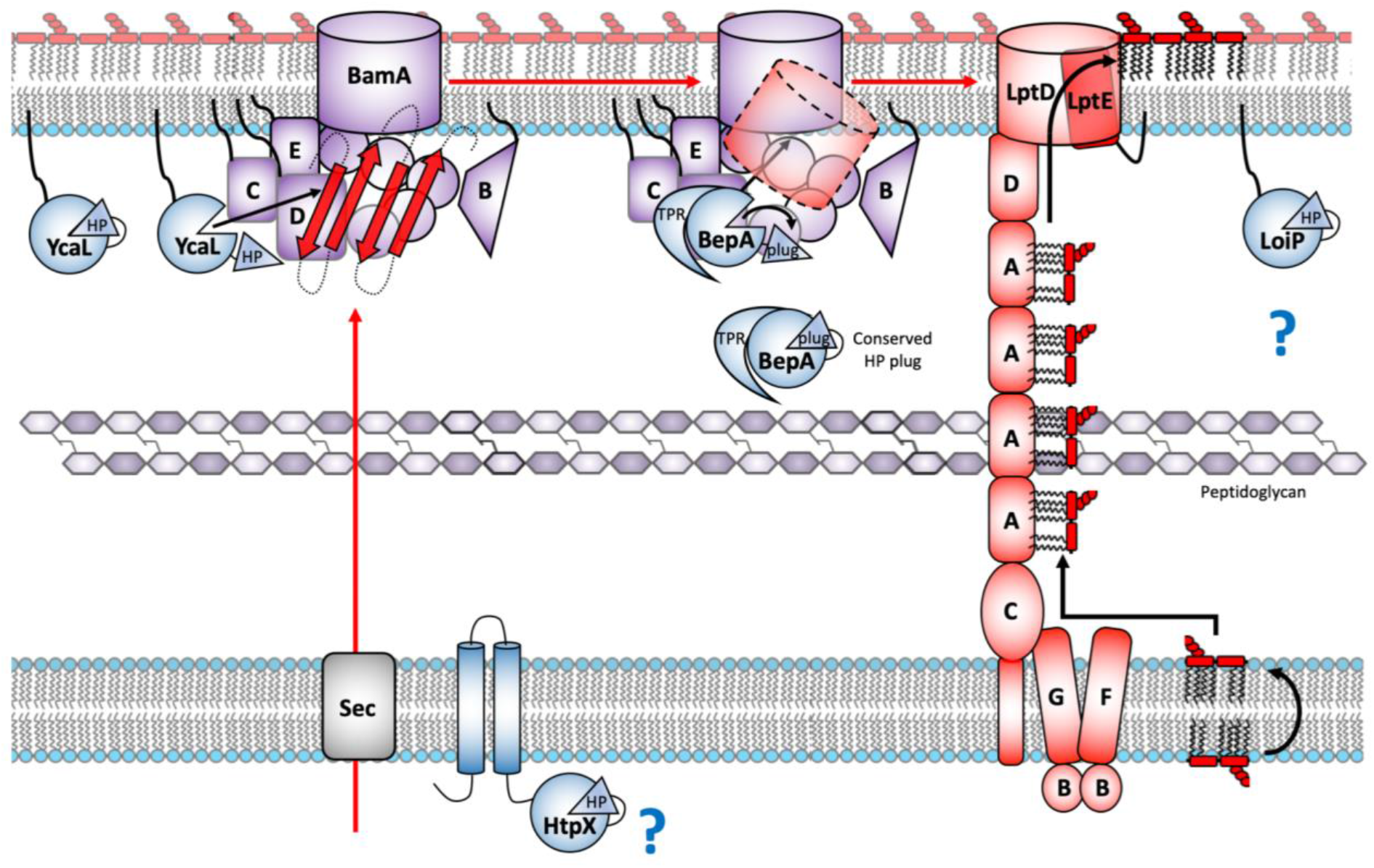
Regulation of the stages of membrane protein biogenesis by the *E. coli* HP-plug M48 metalloproteases. Model for the proteolytic quality control of different stages in integral membrane protein biogenesis by the four *E. coli* M48 metalloproteases, HtpX, YcaL, BepA and LoiP, each of which contains the conserved regulatory HP active site plug (adapted and updated from Soltes *et al* (22)). HtpX is an IM localised M48 metalloprotease targeting misfolded integral membrane proteins, however the targets remain elusive. YcaL is an OM localised lipoprotein specifically targeting Bam-associated, unfolded, OMPs, whereas BepA is a periplasmic metalloprotease targeting the next stage in OMP biogenesis, Bam-engaged partially-folded β-barrels (22). Lastly, LoiP is another OM localised lipoprotein, however LoiP substrates remain uncharacterised. All four of the *E. coli* M48 metalloproteases encode a conserved regulatory active site plug mechanism and appear to be involved in proteolytic quality control of specific stages in integral membrane protein biogenesis.

We identify specific residues in the pocket and cavity formed by the TPR that are important for function. The TPR cavity has previously been shown to be the site of interaction with the Bam complex (30) and here we demonstrate that specific conserved arginine residues within the cavity are important for function and for BepA-mediated degradation of BamA under conditions of stress. While this shows that these residues within TPR are important for BepA function, and potentially substrate recognition, this is not illuminating for the substrate recognition mechanisms of the three remaining *E. coli* M48 metalloproteases. YcaL and LoiP are OM lipoproteins that lack the TPR (22, 44), therefore they do not contain the key residues identified here in the BAM complex interaction cavity or the negatively charged ditch that we demonstrate as important for function. YcaL has been shown to target BAM-engaged substrates that have yet to fold (22). However, considering that it lacks the TPR domain, which is required for BepA interaction with the BAM complex (30), it must recognize the complex and the stalled substrate through a different mechanism. While the other *E. coli* M48 metalloproteases lack the TPR domain they do all contain the active site plug. This will require further study, but we anticipate that our characterization of the BepA active site plug will be of value for further study of the remaining Gram-negative M48 metalloproteases and indeed for proteins in a wide-range of other organisms.

## Methods

### Expression and purification of BepA

The BepA open reading frame, including N-terminal signal peptide, was codon optimized for expression in *E. coli* and cloned into the IPTG inducible vector pET20b fused to a C-terminal His6-tag (a service provided by Genscript). This vector was transformed into *E. coli* DE3 cells and used for recombinant protein production. Briefly, overnight cultures grown in LB media at 37°C were used as the inoculum for autoinduction media supplemented with 10 μM ZnCl2. The resulting cultures were grown at 37°C to an OD_600_ of ~0.8 before the temperature was changed to 18°C for ~18 hours. Cells were harvested by centrifugation and cell pellets were stored at −80°C.

To purify His-tagged BepA, cell pellets were resuspended in buffer A (20 mM imidazole, pH 7.5; 400 mM NaCl) supplemented with 0.05% Tween20 and lysed by sonication. Cell lysates were clarified by ultra-centrifugation and then incubated with Super Ni-NTA agarose resin (Generon) at 4°C with gentle agitation overnight. The incubation mixture was centrifuged briefly, the supernatant was removed, and the resin was resuspended in buffer A before being loaded onto a gravity-flow purification column. The resin was washed extensively with buffer A, then with 20 ml of Buffer A supplemented with 50 mM imidazole, before washing with buffer B (400 mM imidazole, pH 7.5; 400 mM NaCl; 2 % glycerol). BepA protein, eluted in buffer B, was dialyzed against buffer C (20 mM MES, pH 6.5; 5 mM EDTA) at 18°C for 6 h (to remove metals co-purified with BepA protein) and then dialyzed, extensively with sequential buffer changes, against buffer D (as buffer C but lacking EDTA and instead supplemented with 10 μM ZnCl2 and 150 mM LiSO4) at 18°C. BepA protein was concentrated to ~60 mg/ml by ultra-filtration and then further purified on a HiLoad Superdex 200 26/600 column (GE Healthcare) equilibrated in buffer D. Fractions containing pure BepA protein were pooled and concentrated to 35 mg/ml for use in crystallization trials.

### Crystallization and determination of BepA structure

Purified recombinant BepA was used with proprietary crystal screens (supplied by Molecular Dimensions and Jena Bioscience) in sitting drop crystallization experiments using 2 μl of protein solution and 2 μl of crystallization mother liquor at 18°C. Large crystals were obtained in 0.1 M Na HEPES, pH 7.0, and 8% w/v PEG 8,000 and grew within 30 days. Crystals were cryo-protected by step-wise addition of mother liquor supplemented with 25 % ethylene glycol prior to flash freezing in liquid nitrogen.

Protein crystals were used in X-ray diffraction experiments at the Diamond Light Source synchrotron facility (Oxford, UK). Data for SAD experimental phasing was collected at a wavelength of 1.28 Å and was processed using XDS. A single atom of Zn^2+^ (co-purified with BepA) was identified using SHELXD. This initial map was used for auto-building with Phenix. Models were improved by iterations of refinement using Phenix and manual manipulations in COOT.

### Conservation analysis

The consurf server was used to analyze conservation of surface residues. A multiple alignment of BepA homologues was generated using Clustal Omega and submitted to the consurf server along with the BepA pdb file as a basis for conservation analysis. Further to this, the amino acid sequence of the four M48 metalloproteases encoded by *E. coli* were used to generate a multiple alignment by using Clustal Omega and visualized using ESPript 3.0 (http://espript.ibcp.fr) **(Fig S4)**. Lastly, Pfam was used to visualize conservation within the M48 metalloprotease family through use of the HMM logo and the Skylign web server (http://skylign.org.) (46–48).

### *Mutagenesis of* bepA

Mutations in *bepA* were created using a PCR-based site directed mutagenesis approach using the pET20b:: *bepA::6xHis* vector as template. Briefly, *pET20b::bepA::6xHis* vector was used in 18 cycles of PCR using the Phusion polymerase (NEB) as described by the manufacturer, but using complementary primers containing the desired mutation flanked by at least 15-bp of sequence (Table S1). As a negative control, replica reactions were set up and the polymerase omitted. Template DNA was then digested by addition of 20 units (1 μl) *DpnI* restriction enzyme (NEB: R0176S) and incubation at 37°C for 1 h. The reaction mixture was then used to directly transform NEB DH5-α high-efficiency competent cells. Mutations were confirmed by plasmid isolation and Sanger sequencing (Source Bioscience).

### *Functional screening of mutant* bepA

Parent or Δ*bepA* cells were first transformed with the appropriate *pET20b::bepA::6xHis* vector and stored as glycerol stocks at −80°C. Starter cultures were generated by growth overnight (~16 h) at 37°C with aeration in LB broth (10 g/L tryptone; 5 g/L yeast extract; 5 g/L NaCl) supplemented with 100 μg/ml carbenicillin. Cells were normalized to OD_600_ = 1 and then 10-fold serially diluted before 1.5 μl was spotted onto the relevant LB agar plates. Cells were then incubated at 37°C overnight (~16 h) and the plates photographed for record. Cells were screened on LB agar plates supplemented with 100 μg/ml carbenicillin, vancomycin at the stated concentrations and 2 mM TCEP (tris(2-carboxyethyl)phosphine) where stated.

### Western immuno-blotting

To examine the expression of BepA in Δ*bepA* or parent *E. coli,* cells were grown as described for the functional screening of mutant BepA experiments. For analysis of BepA-mediated degradation of BamA, cells were grown for 16 hours at 37°C in M9 minimal media, supplemented with 0.1% casamino acids, 0.4% glucose, 2 mM MgSO_4_, but with CaCl_2_ omitted. The OD_600_ of the cultures was recorded and cells were isolated by centrifugation then resuspended in Laemmli buffer so that the number of cells in each sample was equivalent. Samples were boiled for 10 min, followed by a brief centrifugation step before being resolved by SDS-PAGE and subjected to western blotting using anti-6xhis antibodies (TaKaRa: 631212), or anti-BamA POTRA antiserum (36), as primary antibody and HRP conjugated anti-rabbit (Sigma Aldrich: A6154) antibodies as secondary for detection by use of the ECL system. Samples were loaded in duplicate and subjected to SDS-PAGE simultaneously, followed by coomassie staining and visualization.

### CPRG permeability assay

Following double transformation with the relevant *pET20b::bepA::6xHis* plasmid and the pRW50/CC-61.5 *lac* reporter plasmid (35), cells were grown to mid-exponential phase (OD_600_ 0.4-0.6) in LB broth with aeration and harvested by centrifugation. Cells were resuspended in LB broth to an OD_600_ of 0.1 and 5 μl used to inoculate 96-well culture plates containing 150 μl LB agar supplemented with CPRG (Chlorophenol red-β-D-galactopyranoside – Sigma) (20 μg/ml), carbenicillin (100 μg/ml) and tetracycline (15 μg/ml). 96-well plates were incubated at 30°C and the optical density 300-800 nm monitored every 20 min for 48 h. By using the absorbance of LacZ^-^ strains unable to turn over CPRG, we created an estimating function that predicts the expected absorbance due to cell growth at 575 nm (CPR peak absorbance) using the absorbance at 450 nm and 650 nm. By subtracting the actual absorbance at 575 nm, from the expected growth-related absorbance we derive the CPRG turnover score, which is exclusive to cell membrane permeability. For both expected and measured absorbance at 575 nm, the timepoint of 24 h post-inoculation was used.

### LPS labelling, Lipid A isolation and analysis

Labelling of LPS, Lipid A purification, TLC analysis and quantification were done exactly as described previously (42). Briefly, starter cultures were incubated at 37°C overnight with aeration in LB broth supplemented with 100 μg/ml carbenicillin. Starter cultures were then subcultured into 5 ml LB broth supplemented with 100 μg/ml carbenicillin and the experiment completed precisely as described previously including the addition of the positive control, in which the parent strain was exposed to 25 mM EDTA for 10 min prior to harvest of cells by centrifugation in order to induce PagP mediated palmitoylation of Lipid A (42). Experiments were completed in triplicate and the data generated was analyzed as described previously.

## Data Availability

The BepA X-ray structure has been deposited in the PDB with the accession number 6SAR.

## Acknowledgements

We thank Professor Jeff Cole for stimulating discussion and assistance with manuscript editing. This work was funded by BBSRC and MRC grants to IRH. The lipid A palmitoylation work was supported by the Singapore Ministry of Education Academic Research Fund Tier 2 (MOE2013-T2-1-148) grant (to SSC).

## Author contributions

BepA was identified as a target for study by FM and the project was facilitated by IRH. The BepA mutant was made by FM. Production of the protein was completed by YS, whereas further production, purification, crystallization and modelling of the BepA structure was done by ITC and ALL. Consurf analysis, mutagenesis, western blotting and functional analysis screens were done by JAB with guidance from ALL. Lipid A palmitoylation assays were done by JAB and ZSC under the supervision of SSC. CPRG permeability assays were also completed by JAB with the advice and supervision of MB. CPRG assay data was processed and analyzed by MB and GK. Manuscript preparation was completed by JAB and general project design was done by JAB with the guidance of ALL and IRH. JAB, IRH, MB, ITC and ALL contributed to manuscript editing.

## Supporting information

**Table S1.**
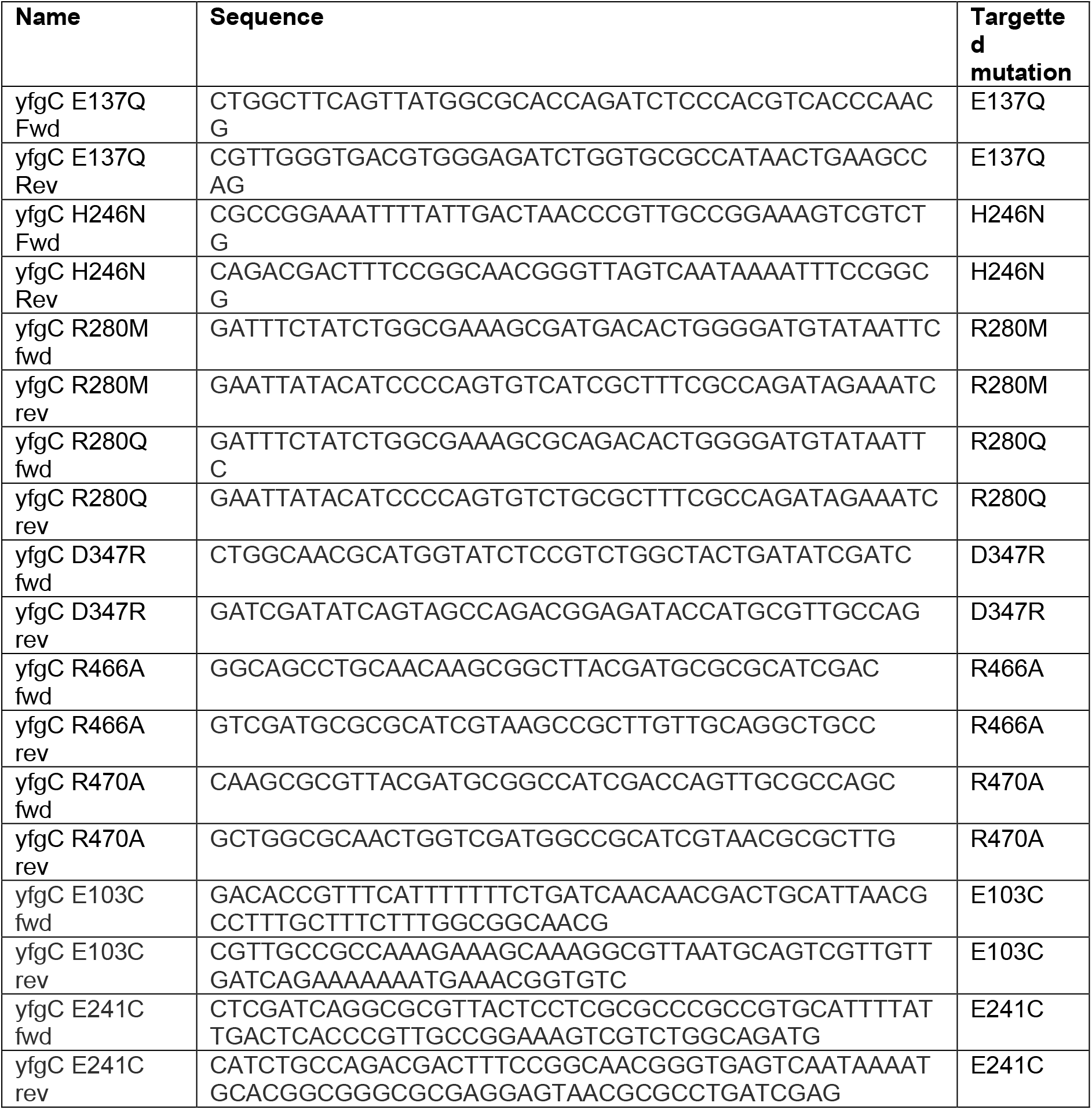
Oligonucleotides used in this study

**Figure S1.**
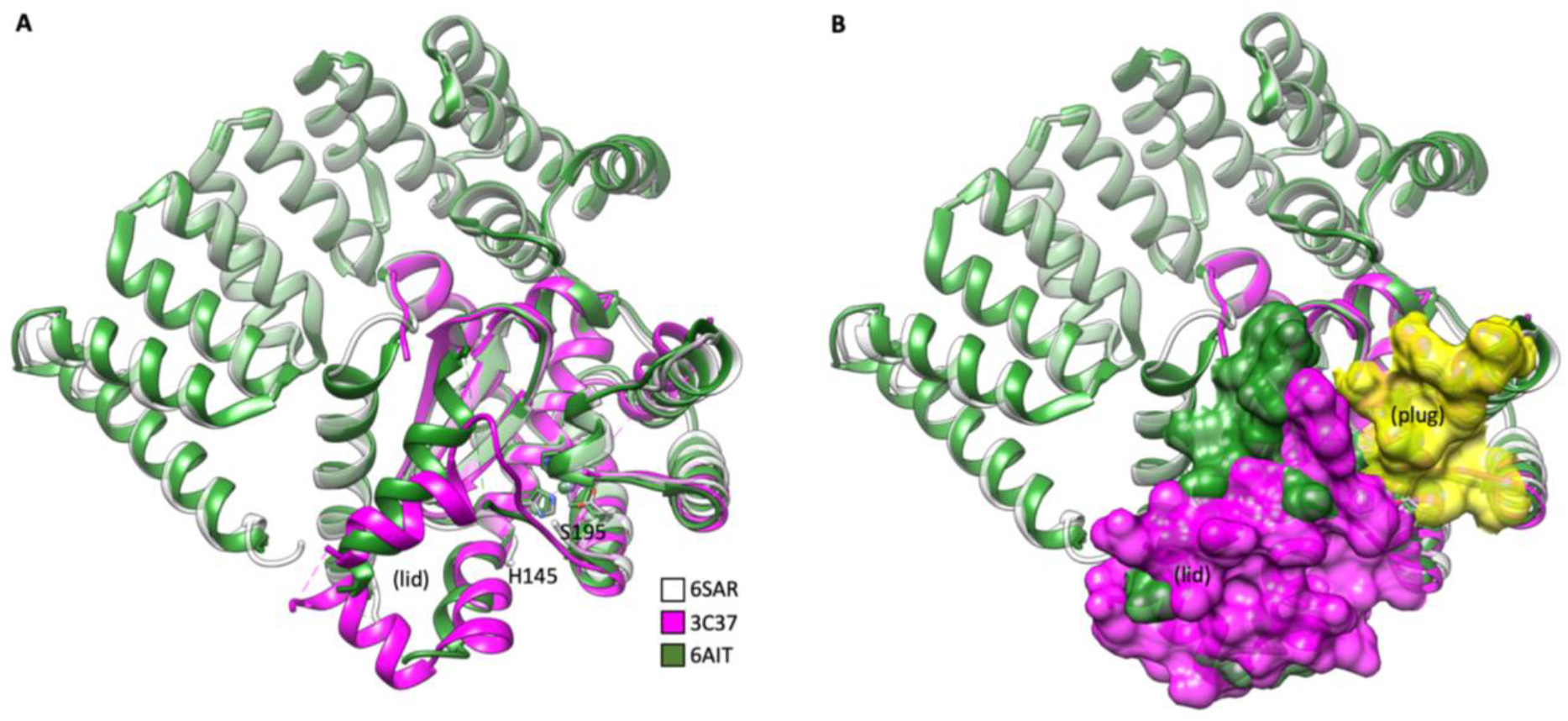
Comparison of PDB: 3C37 and BepA shows the active site lid. Comparison of the *Geobacter sulfureducens* M48 protease structure (PDB: 3C37 – magenta) with that of the BepA structure presented here (PDB: 6SAR – white) and that presented previously (6AIT – green) shows the active site lid formed by the missing residues H145-S195. **A.** Alignment of 6SAR, 6AIT and 3C37 as ribbon representation **B.** Alignment of 6SAR, 6AIT and 3C37 as ribbon representation with surface representation shown for the active site plug of 6SAR (yellow) and the active site lid of 3C37 (magenta) and 6AIT (green) to demonstrate occlusion of the BepA active site.

**Figure S2.**
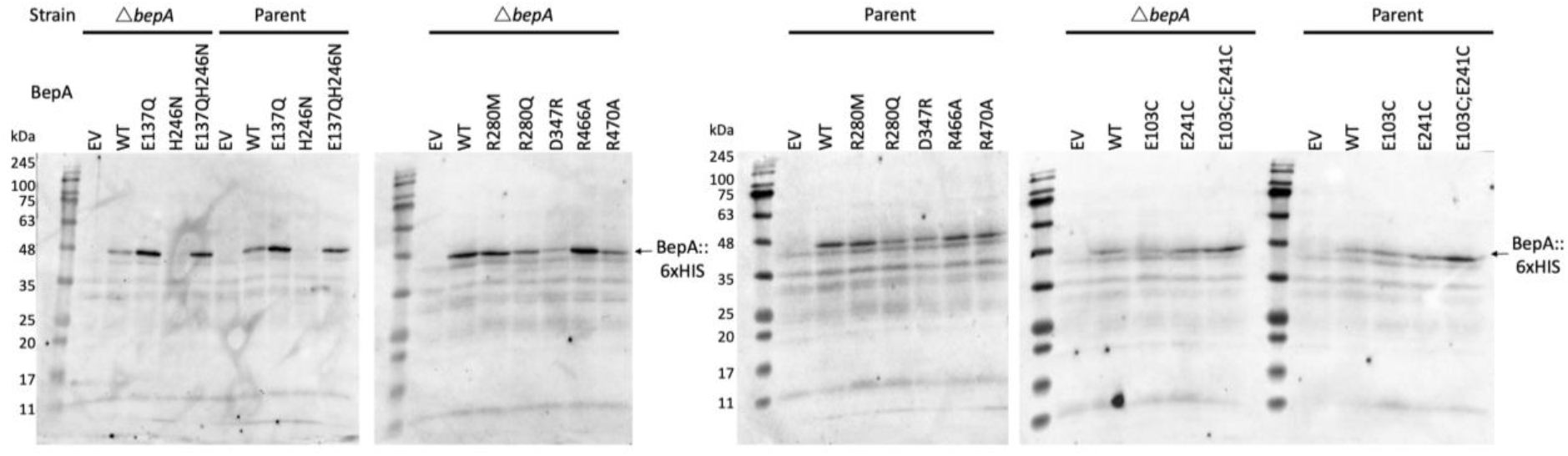
Western immuno-blotting analysis of BepA::6xHis expression. Analysis of BepA expression by western immuno-blotting analysis. Cells carrying pET20b encoding WT or mutated copies of BepA in the parent or Δ*bepA* strain background were harvested and resuspended in Laemmli buffer so that the number of cells in each sample was equal. Following a brief centrifugatiuon step, proteins were separated by SDS-PAGE and transferred by western blot. The empty vector control is labelled EV. Western blotting was completed using anti-6xHis antibody raised in mice and anti-mouse::HRP secondary antibody to target the BepA::6xHis protein in samples used for vancomycin sensitivity screens.

**Figure S3.**
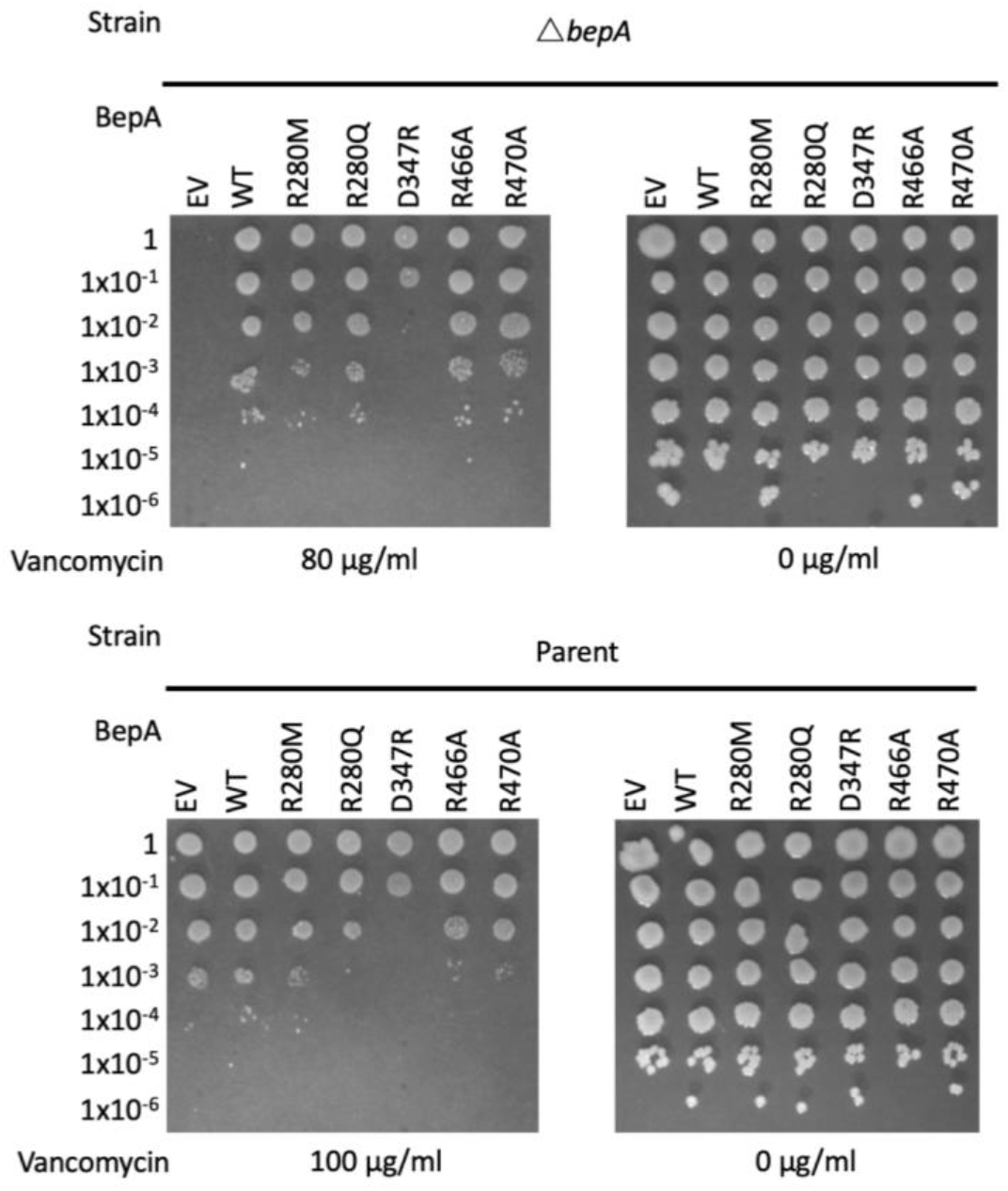
Vancomycin sensitivity screen for pocket and TPR cavity mutants. Screen for vancomycin sensitivity of cells carrying pET20b encoding WT or mutant copies of BepA in the parent or Δ*bepA* strain background. The empty vector control is labelled EV. Cells are normalised to OD_600_ = 1 and ten-fold serially diluted before being spotted on the LB agar containing the indicated antibiotics (all plates contain 100 μg/ml carbenicilin additionally).

**Figure S4.**
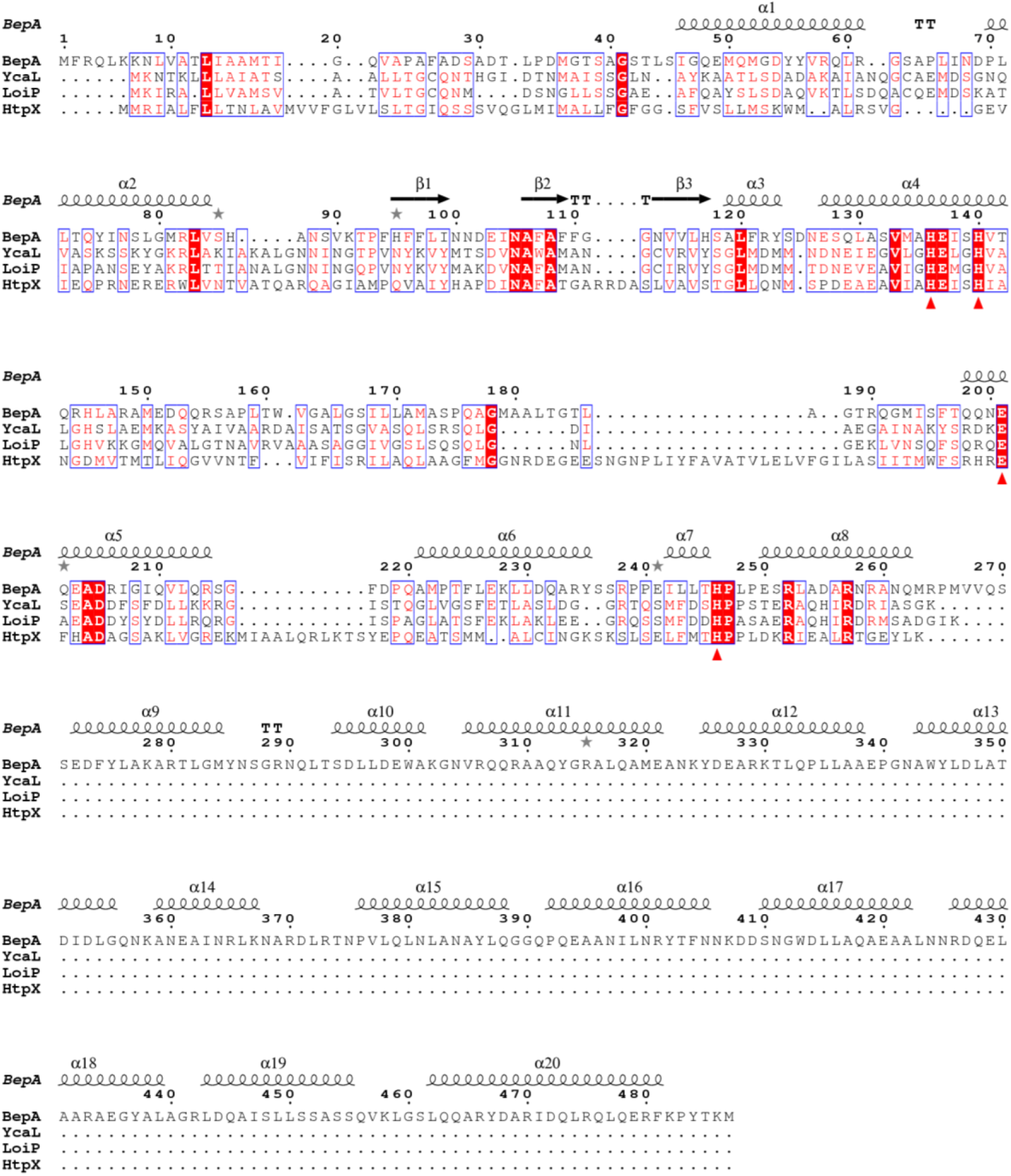
Full multiple alignment of BepA, YcaL, LoiP and HtpX. Amino acid sequences for *E. coli* BepA, YcaL, LoiP and HtpX were submitted to Clustal Omega (https://www.ebi.ac.uk/Tools/msa/clustalo/) in order to generate a multiple alignment to allow analysis of amino acid conservation and subsequently processed using ESPript 3.0 (http://espript.ibcp.fr) (49, 50). Sequences are named on the left and numbered above the line based on the BepA sequence. Gaps in the alignment are represented by dots. A single fully conserved residue is highlighted red and the zinc co-ordinating residues are labelled with a red triangle under the residue. BepA secondary structure is represented on the top line with α-helices labelled with a spiral and β-sheets by arrows.

